# Cross-Dataset Identification of Human Disease-Specific Cell Subtypes Enabled by the Gene Print-based Algorithm--gPRINT

**DOI:** 10.1101/2023.11.05.565588

**Authors:** Ruojin Yan, Chunmei Fan, Shen Gu, Tingzhang Wang, Zi Yin, Xiao CHEN

## Abstract

Despite extensive efforts in developing cell annotation algorithms for single cell RNA sequencing results, most algorithms fail to achieve cross-dataset mapping of cell subtypes due to factors such as batch effects between datasets. This limitation is particularly evident when rapidly annotating disease-specific cell subtypes across multiple datasets. In this study, we present gPRINT, a machine learning tool that utilizes the unique one-dimensional “gene print” expression patterns of individual cells. gPRINT is capable of automatically predicting cell types and annotating disease-specific cell subtypes. The development of gPRINT involved curation and harmonization of public datasets, algorithm validation within and across datasets, and the annotation of disease-specific fibroblast subtypes across various disease subgroups and datasets. Additionally, we created a preliminary single-cell atlas of human tendinopathy fibroblasts and successfully achieved automatic prediction of disease-specific cell subtypes in tendon disease. Furthermore, we conducted an exploration of key targets and related drugs specific to this subtype in tendon disease. The proposed approach offers an automated and unified method for identifying disease-specific cell subtypes across datasets, serving as a valuable reference for annotating fibroblast-specific subtypes in different disease states and facilitating the exploration of therapeutic targets in tendon disease.

## Main

Disease-specific cell subtypes refer to subgroups of cells that exhibit distinct characteristics and functions within a particular disease context. Identifying these cells has profound implications for understanding the pathogenesis, diagnosis, treatment, and prevention of diseases. For example, in-depth exploration of disease-specific cell subtypes can lead to a better understanding of the pathogenesis and progression of diseases and facilitating the development of targeted therapeutic approaches for specific cellular subtypes, such as the study of disease-specific cell subpopulations and molecules in chronic rhinosinusitis (CRS), which identifies potential therapeutic approaches to treat CRS^[1]^, and the study of the correlation between intertissue cell subtype structure of fibroblasts and different diseases, which provides information for treatment approaches^[2]^. In addition, studying disease-specific cell subtypes can provide critical information for repurposing existing drugs and developing new drugs to conduct clinical trials^[3]^.

Currently, the definition and classification of disease-specific cell subtypes are mostly accomplished manually^[4][5]^. Due to the current lack of clear standards and consensus on the definition of cell subtypes^[6]^, manual annotation results are inconsistent and incomparable, thus limiting the research progress and clinical application of disease-specific cell subtypes. Given the complexity and heterogeneity of diseases, accurately and uniformly defining and identifying disease-specific cell subtypes is essential. Many efforts have been made in developing cell automatic annotation algorithms. For example, scDeepsort^[6]^ and SingleR^[1]^ can annotate cells to the cell type level based on large-scale cell information databases such as HCL^[7]^ and MCA^[8]^; scType^[9]^, CellAssign^[10]^ and SCSA^[11]^ can annotate cells based on existing marker gene databases; scmap^[3]^ and Cellid^[12]^ can map cell types between different data sets. However, the above algorithms still face challenges in accurately annotating and verifying cell subtypes, which are more refined than cell types, across different data sets, mainly due to the absence of marker genes for cell subtypes and the presence of batch effects^[1][3][6]-[13]^. However, there is currently a lack of algorithm which is precise and unified annotation of disease-specific cell subtypes across datasets.

To address these challenges, we present Generalized PlatfoRm for cell type Identification across diverse single-cell RNA-Seq experiments using Neural neTworks (gPRINT), an R and Python based package, to accomplish unified mapping of disease-specific cell subtypes, which may have different annotation hierarchies, obtained from any tissue .The gPRINT is based on the principle of sound recognition in noisy environments^[14]^. Firstly, we reordered the genes of each cell according to their positions in the standard Human Reference Genome Build 38 (HG38), and then ploted the levels of gene expression to obtain the “gene print” of each cell. Next, we used the neural network to reduce the impact of “background noise” in single-cell analysis^[15]^, ultimately achieving rapid and uniform annotation of disease-specific cell subtypes.

In order to prove the ability of gPRINT in cell type and subtype prediction, we first explored the feasibility and scalability on multiple publicly available scRNA-seq data sets and the HCL database^[7]^ (Figure 1). These data sets include 26 tissues including human pancreas tissue^[19–24]^, colorectal tumors^[25]^, brain tissue^[26]^, bladder, bone marrow, cervix, esophagus, heart and ileum^[7]^ with a total of 159,302 cells (Fig 1A). Subsequently, we demonstrated the ability of gPRINT to predict cell identity across levels and time on the human skeletal muscle atlas^[27]^. Finally, cell subtype identity mapping across disease subgroups was completed on the fibroblast atlas under human NSCLC disease^[28]^, with an average accuracy rate of 84.30%. At the same time, based on this atlas data, the unified annotation of fibroblast subtypes under NSCLC disease across data sets^[2]^ was completed. These results demonstrate the feasibility and accuracy of the gPRINT algorithm in accurately and uniformly annotating disease-specific cell subtypes across data sets. After unified annotation, the disease mechanism can be analyzed for this specific subtype, key treatment targets for different diseases can be found, and old drugs can be screened or new drugs can be developed, providing an important foundation for precision medicine.

**Figure 1.**
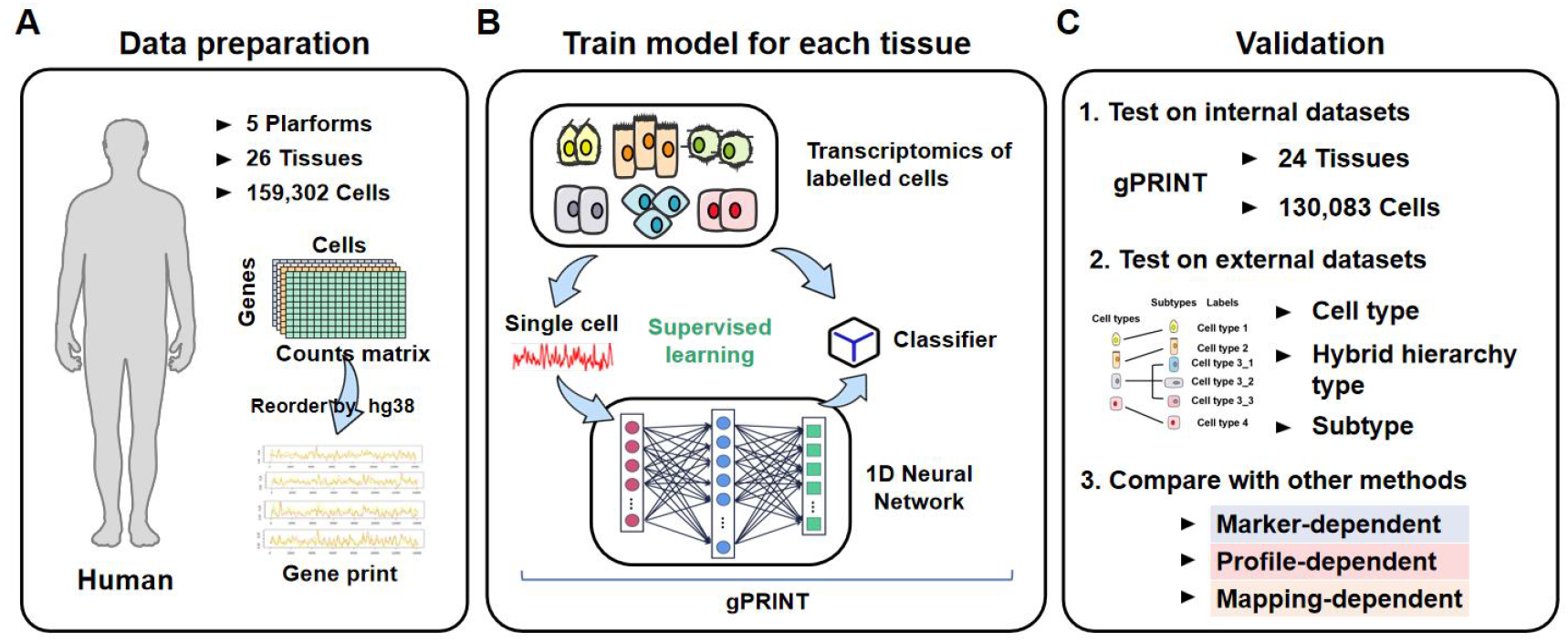
General conceptual framework and validation of gPRINT. (A) Selected human single-cell transcriptomic profiles from HCL and other public datasets for training and validating gPRINT. Human single cell data includes 159,302 cells from 26 tissues and 5 platforms. (B) For each cell, a neural network was constructed of this cell and its “gene print” for supervised learning gPRINT with known cell labels from the transcriptomic atlases for each tissue. (C) The performance of gPRINT was tested using an internal human dataset and an external test dataset containing single-cell transcriptomic data from multiple tissues. For the human external test data set, it was verified at the cell type, mixed cell type and subtype, and cell subtype levels. gPRINT was also compared with other different types of annotation algorithms on various datasets.

## Results

### 2.1 gPRINT: A Tool for Unified Annotation of Disease-Specific Cell Subtypes Based on Gene Fingerprints

To address the challenge of accurately annotating cell identities within complex annotation hierarchies and enabling unified identification of disease-specific subtypes, we developed gPRINT. It is a software package based on R and Python that enables the automatic prediction of cell identities in highly complex annotation hierarchies obtained from any tissue or sequencing platform. Just as each individual possesses unique fingerprints and voiceprints^[29]^, each cell has its own “gene print”. The gPRINT algorithm accomplishes this by reordering the genes expressed in each cell according to the human reference genome sequence HG38 (refer to the “Methods” section) and plotting the gene expression levels to generate its unique “gene print” (Figure S1). Building on the principles of deep learning applied in voice recognition, the algorithm treats the positional information of gene open expression as temporal information in a sound wave. Each gene interval is treated as a frame segment in a sound wave, and a one-dimensional neural network is used to learn from a specific reference dataset and automatically predict cell identities in the query dataset. Notably, this approach includes the additional feature of gene-to-gene positional information (Figure 1B).In summary, gPRINT algorithm uses the similarity between single-cell gene expression profiles and reference expression profiles to complete the identity mapping of single cells at the cell subtype level in a scalable manner (Figure S1, refer to the “Methods” section). The gPRINT_group algorithm, which relies on one-dimensional backpropagation (BP) neural networks^[30]^, learns from the “feature prints” derived from marker genes for each cell type in the reference dataset. It then performs cell identity recognition on the query dataset based on cell clusters (Figure S1, please refer to the “Methods” section).

In summary, this algorithm empowers users to create trained models based on reference scRNA-seq datasets obtained from any human tissue or sequencing platform. These models can then be used to predict cell types for new datasets featuring similar cell types and subtypes.

### 2.2 Internal Test Sets for Identifying Cell Type Recognition

To verify the feasibility of gPRINT for cell type annotation, we first used a classic pancreas dataset in the field of cell annotation that has been annotated and validated by different algorithms^[19]^. In this study, we employed five well-established pancreatic datasets (Wang^[20]^, Xin^[21]^, Muraro^[22]^, Baron^[23]^ and Segerstolpe^[24]^) to validate the algorithm.

To evaluate the classification performance of gPRINT, we first converted the cellular gene expression matrix of each data set into a “gene print” expression form, and then randomly divided the data into 70% training partitions and 30% testing partitions. We performed 5-fold cross-validation on the 30% partition, involving independently retraining the model five times and testing it on randomly partitioned data. We assessed four key performance metrics, including model accuracy, precision, recall, and F1 score (Figure 2A, refer to the “Methods” section). For cell annotation in the Baron dataset^[23]^, the model demonstrated high overall accuracy, achieving approximately 98.4% across 5 independent rounds of training (Figure 3). The model also exhibited a high degree of sensitivity, with a recall value of 98.4% ± 0.30%, and maintained an overall precision score of 98.2% ± 0.4%. Similarly, gPRINT’s performance on other pancreatic datasets was impressive, with an average accuracy of 97.8% on the Muraro dataset^[22]^, 83.4% on the Segerstolpe dataset^[24]^, 87.6% on the Wang dataset^[20]^, and 92.6% on the Xin dataset^[21]^ (Figure 2B). These results collectively demonstrate the feasibility of the gPRINT algorithm for highly accurate cell identity prediction. Taking the Muraro dataset^[22]^ as an example, 5-fold cross-validation on the 30% test set confirmed highly accurate predictions. All cell categories had prediction accuracies exceeding 93%, with four categories such as acinar and alpha cells achieving prediction accuracies exceeding 99%. Confusion matrices for other datasets are shown in Supplementary Figure S2.

**Figure 2.**
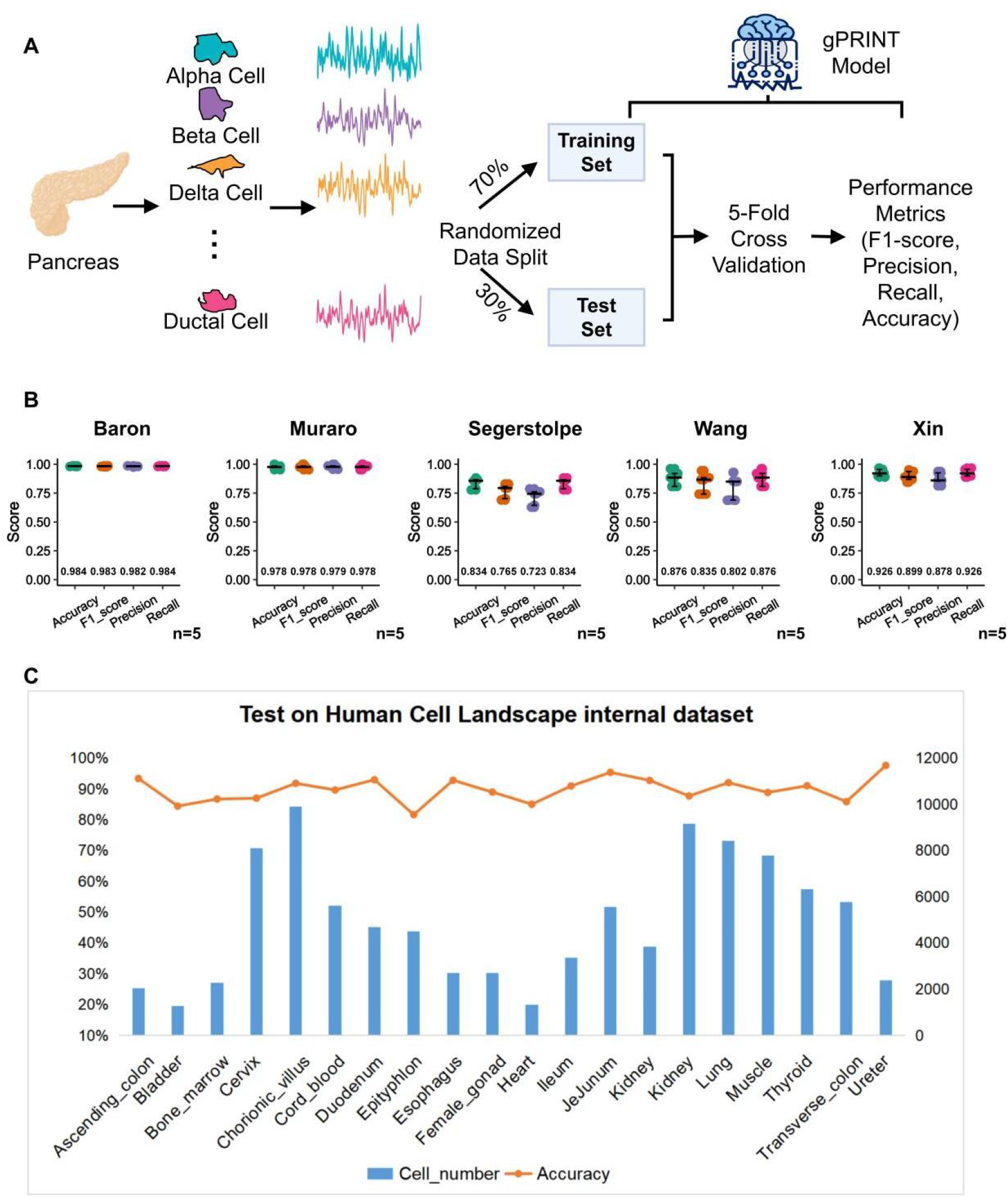
Results of internal validation of gPRINT on small public datasets and the HCL database to determine the classification performance of the algorithm. (A) Demonstration of the gPRINT annotation scheme within pancreatic tissue. All “gene prints” were randomly split into 70% as a training set and 30% as a test set, and then 5-fold cross-validation was used for final testing of model accuracy. Specific evaluation indicators include F1-score, Precision, Recall, and Accuracy. (B) Performance measures of 5-fold cross-validation for each dataset annotation. Error bars represent 95% confidence intervals. Test independent folding for n = 5. (C) Classification accuracy of gPRINT internal annotations on the HCL database. The left side of the coordinate axis represents the accuracy rate, and the right side represents the number of cells.

**Figure 3.**
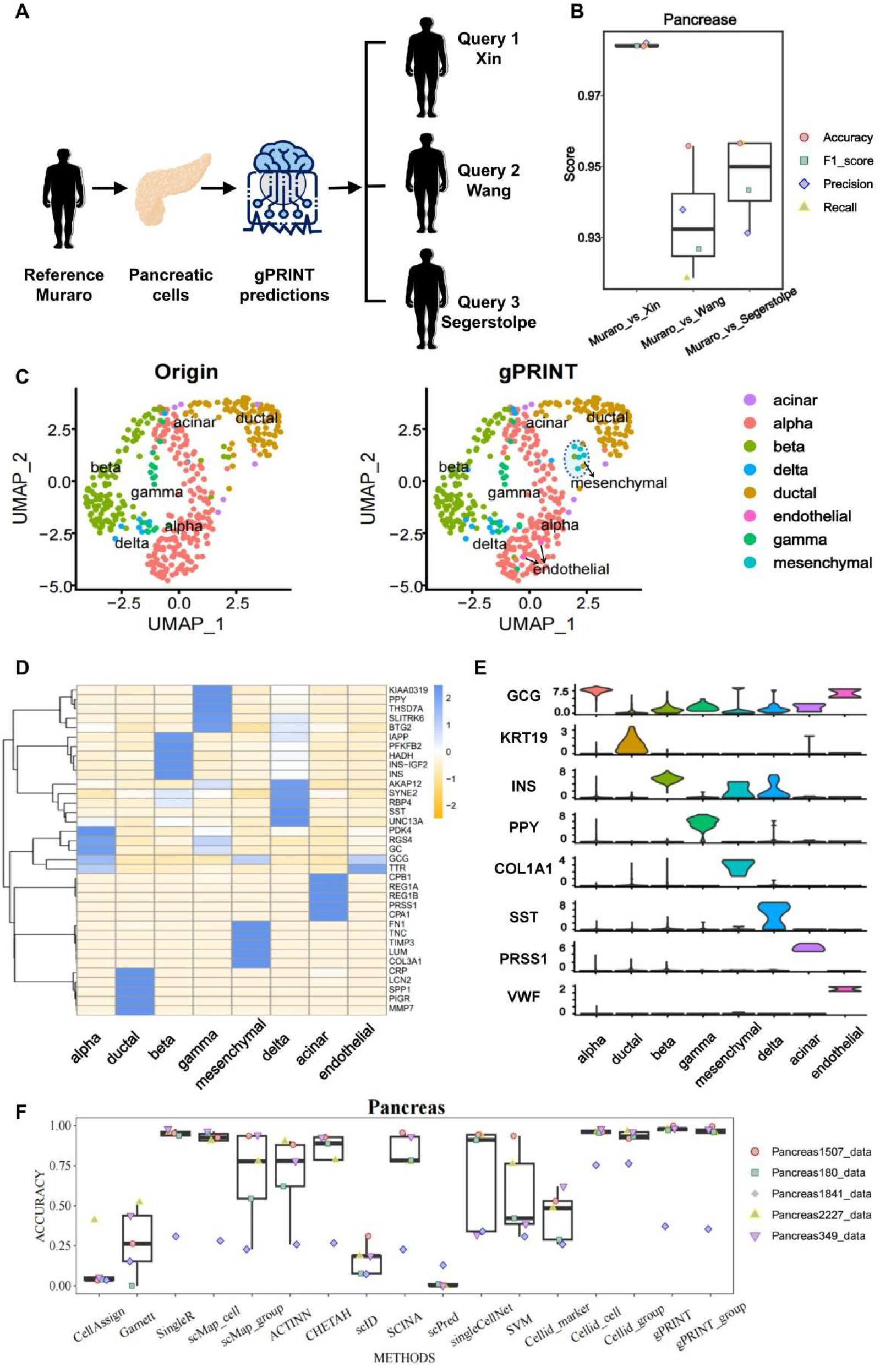
Performance of gPRINT cell type identity mapping from different scRNA-seq datasets within pancreatic tissue. (A) Demonstration of the gPRINT cross-tissue annotation scheme within pancreatic tissue. Using the Muraro data set as a reference, complete the annotation of the Xin, Wang and Segerstolpe data sets (B) Evaluation indicator bar chart of the annotation results of different data sets. Boxplots summarize method’s F1 scores, and show the median (center line), interquartile range (hinges) and 1.5 times the interquartile range (whiskers). (C) UMAP plot display of cells from the Wang dataset. The left side shows the cell types defined in the original text, and the right side shows the results of cell type prediction based on gPRINT, where the colors of the cells represent different cell types. (D) Average of the top 5 differentially expressed markers across all gPRINT-predicted cell types and presented as a heatmap. (E) Violin plot representing expression levels of marker genes across all different cell types. (F) Boxplots summarize method’s F1 scores, and show the median (center line), interquartile range (hinges) and 1.5 times the interquartile range (whiskers).

We conducted another internal validations on single-cell datasets from human colorectal tumors (Li dataset^[25]^) and human brain tissues (Camp dataset^[26]^), resulting in accuracy rates exceeding 92% (Figure 2C). Furtherly, we also performed internal validations on one of the biggest human cell landscape (HCL) database^[7]^, including the bladder, bone marrow, cervix, esophagus, heart, and ileum (Figure 2C), with accuracy rates exceeding 80%. These validations collectively demonstrate the feasibility of gPRINT across various tissues and datasets.

### 2.3 Cross-Datasets Cell Type Identity Mapping within the Same Tissue

From the results mentioned above, it’s evident that gPRINT demonstrates a degree of feasibility and accuracy when transferring labels from one dataset to another. The accuracy of gPRINT in cross-dataset label mapping indicates that the algorithm, to a certain extent, mitigates the impact of batch effects when annotating different small datasets. This can be attributed to the use of a 1D CNN model in our approach, where the convolutional layers act as filters. The multi-layer convolutional and pooling layers enable the algorithm to extract the most relevant features from one-dimensional expression data. This ability allows it to filter out noise introduced by batch effects and reduce the influence of batch effects on cell annotation extrapolation. Furthermore, since the algorithm learns from nearly all expressed genes and captures relevant features through the convolutional layers, it theoretically makes optimal use of the gene information even in low-depth and low-quality single-cell data. This results in more accurate cell annotation information and avoids the problem of low accuracy in cell annotation that arises from the use of marker gene-based annotation methods, which are limited by missing gene expression data for many genes.

Specifically, using the Muraro dataset^[22]^ as a reference and annotating the Wang^[20]^ dataset as an example, we successfully obtained annotations for cell types such as acinar, beta, gamma, alpha, delta, and ductal using gPRINT. Additionally, we were able to annotate cell types such as mesenchymal and endothelial, which had not been previously defined in Wang’s study^[20]^ (Figure 3C). We further validated the annotations assigned in the map construction by performing differential gene expression analysis on all major annotated cell types and confirmed the unique expression of the major cellular markers reported in the 8 major cell populations identified (Figure 3D). Subsequently, violin plots were generated for subtypes using marker genes to illustrate the re-annotation of cell types by gPRINT (Figure 3E). These plots revealed high expression of mesenchymal and endothelial marker genes in the re-annotated cell subtypes, affirming the correctness of gPRINT’s annotations. The algorithm effectively corrected the cell types in the original dataset, achieving cross-data set annotation.

To further assess the accuracy and scalability of gPRINT, we performed annotations on five low-quality pancreatic datasets (Figure 3F, S3). The average accuracy achieved was above 97.22% (matrix data) and 95.51% (marker genes) or higher. Compared to other algorithms such as scmap^[3]^, CellAssign^[10]^, Garnett^[24]^, SingleR^[1]^, ACTINN^[32]^, CHETAH^[33]^, scID^[34]^, SCINA^[35]^, scPred^[36]^, SVM^[30]^, and Cellid^[12]^, gPRINT demonstrated superior performance.

For datasets where all algorithms could annotate correctly, including human_Pancreas349, human_Pancreas2227, human_Pancreas180, and human_Pancreas1507^[6]^, gPRNIT and gPRINT_group consistently achieved high accuracy (>97% and >95%) across all reference-to-query cell-type assignments. In the case of the human_Pancreas1841^[6]^ dataset, which none of the algorithms could accurately annotate, gPRINT also did not give correct results for the annotations. This suggests that gPRINT avoids overfitting, and the dataset itself may have incorrect annotations. Among the tested algorithms, CellAssign^[10]^, Garnett^[24]^, and SCINA^[35]^ are cell annotation algorithms based on marker genes, while SingleR^[1]^, scmap^[3]^, ACTINN^[32]^, CHETAH^[33]^, scID^[34]^, scPred^[36]^ and SVM^[30]^ are cell annotation algorithms based on reference datasets. Since the marker genes for various cell types in pancreatic tissue are relatively well-established, and pancreatic single-cell data maps have been constructed on the HCL, both types of cell annotation algorithms can effectively extend their annotations in pancreatic tissue.

**Table 1.**
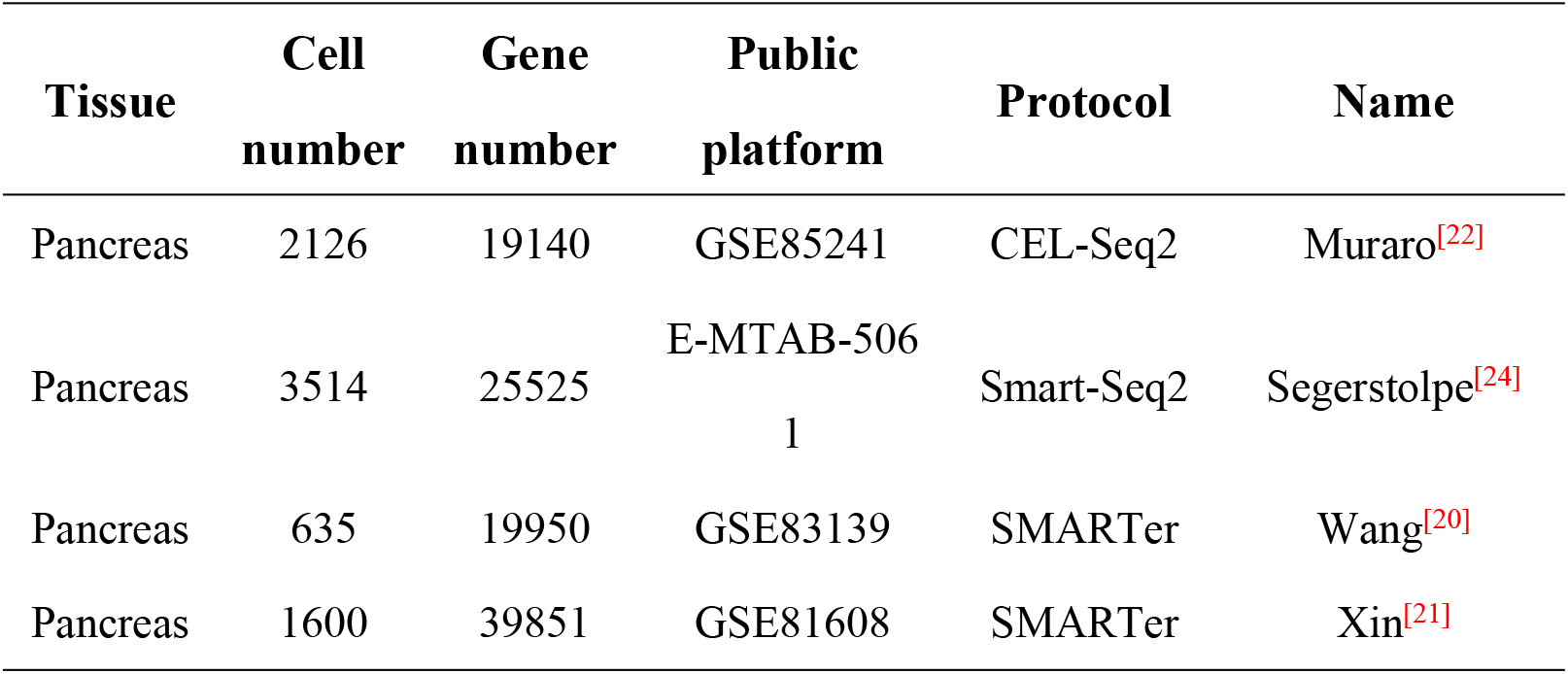
Detailed information of different data sets on pancreatic tissue.

### 2.4 Cross-Time Label Recognition on the Human Skeletal Muscle Atlas Based on Complex Cell Type Hierarchies

We also utilized the gPRINT algorithm to map cell identity across time in the human skeletal muscle atlas dataset GSE147457, which includes manually assigned cell types^[27]^. This allowed us to assess the feasibility of the algorithm in annotating fine-grained fibroblast subtypes and evaluating its advantages compared to other algorithms, particularly in scenarios with mixed cell types and detailed cell labels (Figure 4A). First, we attempted to annotate the Human Skeletal Muscle Atlas datasets^[27]^, including the Fetal Wk12_14 and Fetal Wk17_18 datasets, using methods based on signature gene databases. However, we observed that both the scType^[10]^ and Cellid^[12]^ algorithms failed to provide effective annotations. This failure could be attributed to the absence of signature genes in the gene databases as shown in Figure S4. Subsequently, we also employed the SingleR^[1]^ algorithm for the annotation, which relies on large reference databases. Surprisingly, SingleR also proved ineffective in providing accurate annotations. This may be due to the lack of relevant information about these subtypes within the large reference databases, as depicted in Figure S4.

**Figure 4.**
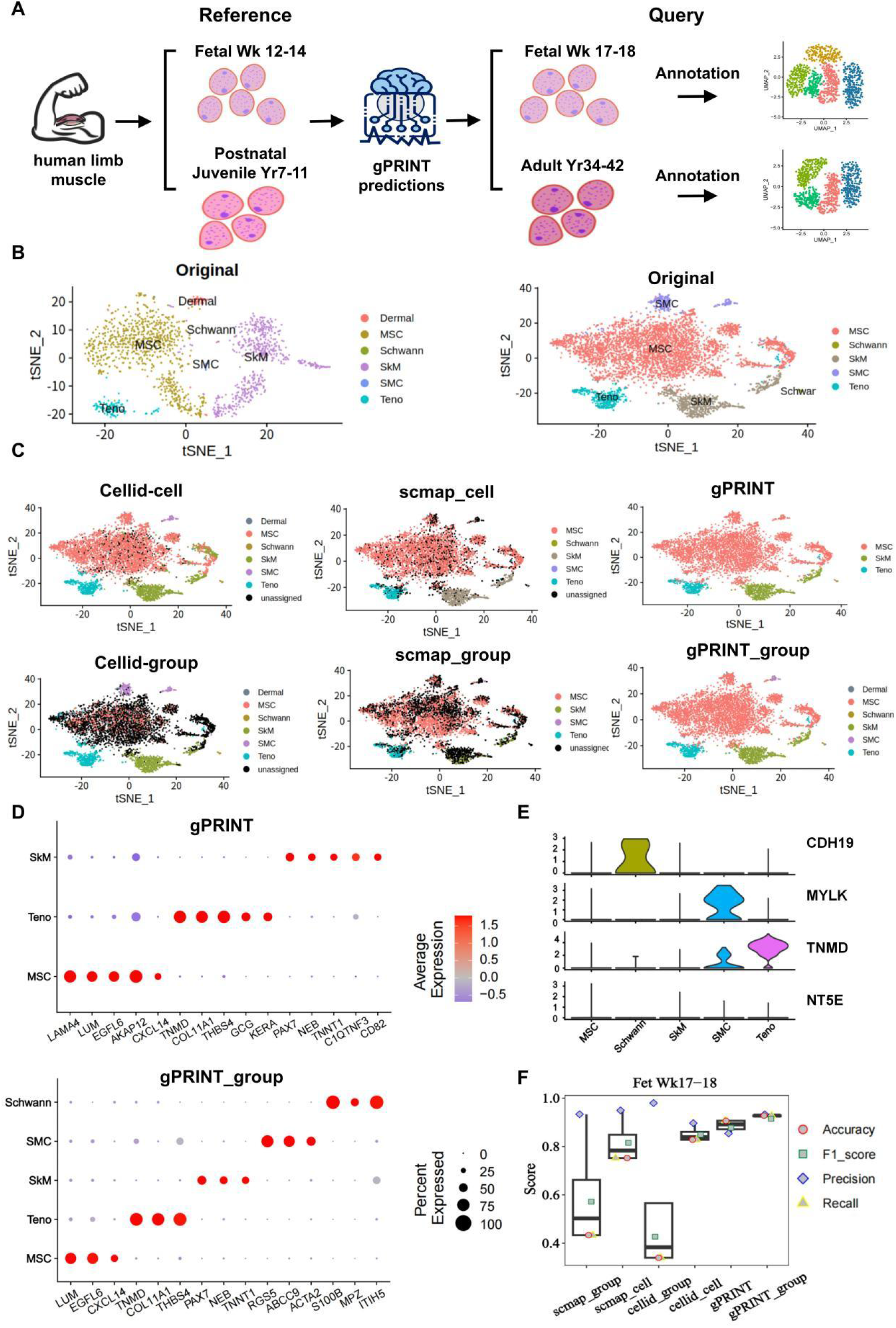
Cell type identity mapping performance of gPRINT on human fetal limb skeletal muscle atlas. (A) Demonstration of the gPRINT annotation scheme across time and samples on the human limb skeletal muscle atlas. On the fetal skeletal muscle atlas, Fetal Wk 12-14 samples are reference data, and Fetal Wk 17-18 samples are test data; on the adult skeletal muscle atlas, Postnatal Juvenile Yr7-11 is reference data, and Adult Yr34-42 samples are test data. (B) t-SNE diagram of cell distribution of different samples on the fetal limb skeletal muscle atlas. The color of the cells represents the cell type. (C) t-SNE plot shows the annotation results of Fetal Wk 17-18 samples by different annotation methods, where the color of the cells represents the cell type. (D) The average of the top 4 or 3 differentially expressed markers in cell types predicted by all gPRINT and gPRINT_group algorithms is presented in the form of a dot plot. (E) Violin plot representing expression levels of marker genes across all different cell types. (F) Bar chart of evaluation metrics of each annotation algorithm on the Fetal Wk 17-18 sample. Boxplots summarize method’s F1 scores, and show the median (center line), interquartile range (hinges) and 1.5 times the interquartile range (whiskers).

Considering the issues mentioned earlier, including the absence of signature genes and the lack of subcellular subtype information in large databases, Then we utilized another two algorithms based on the projecting new samples onto existing reference datasets: scmap^[3]^ and Cellid^[12]^ to perform cross-dataset annotation on the Fet Wk12_14 dataset as a reference for Fet Wk17_18. We found that these two algorithms outperformed methods relying on signature gene libraries or large reference datasets.They could approximately reproduce the cell labels defined in the original study, especially when based on single-cell type learning (Fig4C) .The Fet Wk12_14 dataset included cell labels such as Schwann cells, tenogenic cells (Teno), mesenchymal stem cells (MSC), and muscle cells, with muscle cells comprising smooth muscle cells (SMC) and skeletal muscle cells (SkM) (Figure 4B). However, there were still numerous cells that could not be clearly defined (indicated as black points in the t-SNE plots), and in some cases, smooth muscle cells (SMC) were misclassified as mesenchymal stem cells (MSC). While our algorithm provided clearer results in the annotated t-SNE plots, it was still incapable to accurately label some SMC subtypes and Schwann cell subtypes(Figure 4C). Next, we evaluated the differences between gPRINT and gPRINT_group and the other two algorithms based on specific performance metrics (Figure 4F). Intuitively, through boxplots, swarm plots, and confusion matrices for each cell type, it was evident that gPRINT and gPRINT_group outperformed the other two algorithms in terms of accuracy, recall, and F1-score. Specifically, gPRINT and gPRINT_group consistently achieved high accuracy (>90% and >92%), recall (>90% and >92%), and F1 values (>87% and >91%, respectively), across all evaluated reference-to-query cell-type assignments. The second-best algorithm in this scenario was Cellid_cell, with specific evaluation metrics including accuracy (>82%), recall (>82%), and F1 values (>84%). In this case, the use of the gPRINT_group algorithm proved to be more precise than the gPRINT algorithm. Further examination of violin plots displaying the expression of signature genes for different cell types on gPRINT_group annotated subtypes (Figure 4E) revealed that CDH19, MYLK, TNMD, and NT5E were signature genes for Schwann cells, smooth muscle cells (SMC), tenogenic cells (Teno), and mesenchymal stem cells (MSC)^[31]^, respectively. These marker genes exhibited high expression in the corresponding cell types predicted by the gPRINT_group method, indicating the successful completion of cross-temporal cell type and subtype mapping in the human fetal skeletal muscle atlas from 12-14 weeks to 17-18 weeks.

Next, we proceeded to perform cross-dataset^[27]^ validation on postnatal juvenile (years 7–11) stage and adult (years 34–42) stage datasets from the adult limb skeletal muscle atlas (Figure 4A, S6A). Upon observing the boxplots and swarm plots for evaluation metrics (Figure S6D), it is evident that the gPRINT algorithm outperformed the other algorithms in terms of accuracy, recall, and F1-score. Specifically, gPRINT and gPRINT_group consistently achieved high accuracy (>92% and >93%), recall (>92% and >93%), and F1 values (>91% and >93%, respectively), across all evaluated reference-to-query cell-type assignments. The second-best algorithm in this scenario was Cellid_cell, with specific evaluation metrics including accuracy (>64%), recall (>64%), and F1 values (>74%). Further examination of violin plots displaying the expression of signature genes for different cell types on gPRINT annotated subtypes (Figure S6E) showed that *MYLK*, *SRGN*, and *PDGFRA* were signature genes for SMC, EC-Hema, and FAP cell types^[27]^, respectively. Therefore, we consider our annotation to be correct.

The results above indicate that the gPRINT algorithm can be used to compare datasets from similar biological sources collected by different laboratories. It ensures consistency in annotation and analysis, removes the impact of batch effects on the analysis, and can accurately annotate datasets with mixed cell types and subtypes across different datasets and time points. Hence, we conclude that the gPRINT algorithm can address the challenge of precise mapping of cell subtype labels.

### 2.5 Cross-Dataset Annotation Reveals Disease-Specific Cell Subtypes in Human Fibroblasts within NSCLC Disease Subgroups

Identifying disease-specific cell subtypes has profound implications for understanding the pathogenesis, diagnosis, treatment, and prevention of diseases. And gPRINT algorithm has previously shown its potential to accurately map cell subtype labels, so we further explored the annotation effect of the gPRINT algorithm on cell subtypes under different health and disease subgroups. Due to the differences between healthy and disease states, these differences may mask or interfere with the true differences between cell subtypes, making it difficult to accurately map cell subtypes between healthy and disease subgroups. Similarly, batch effects between different datasets can also interfere with the accuracy of cell subtype mapping.

First, we conducted cell subtype mapping of fibroblasts in the human NSCLC (non-small cell lung cancer) disease atlas in Christopher’s study^[28]^. This was done across healthy and disease subgroups to compare the effectiveness of various algorithms in annotating cell subtypes. This NSCLC atlas, which includes control samples, adenocarcinomas (LUAD), squamous cell carcinomas (LUSC), and large cell carcinomas (LCLC) (Figure 5A,B), employs both multiplex immunohistochemistry and digital cytometry (CIBERSORTx) for the validation of annotated fibroblast cell subtypes (Adventitial, Alveolar, and Myofibroblasts). We utilized this atlas to validate the gPRINT algorithm. We used the Control group as a reference to map the cell subtype labels of the other groups.

**Figure 5.**
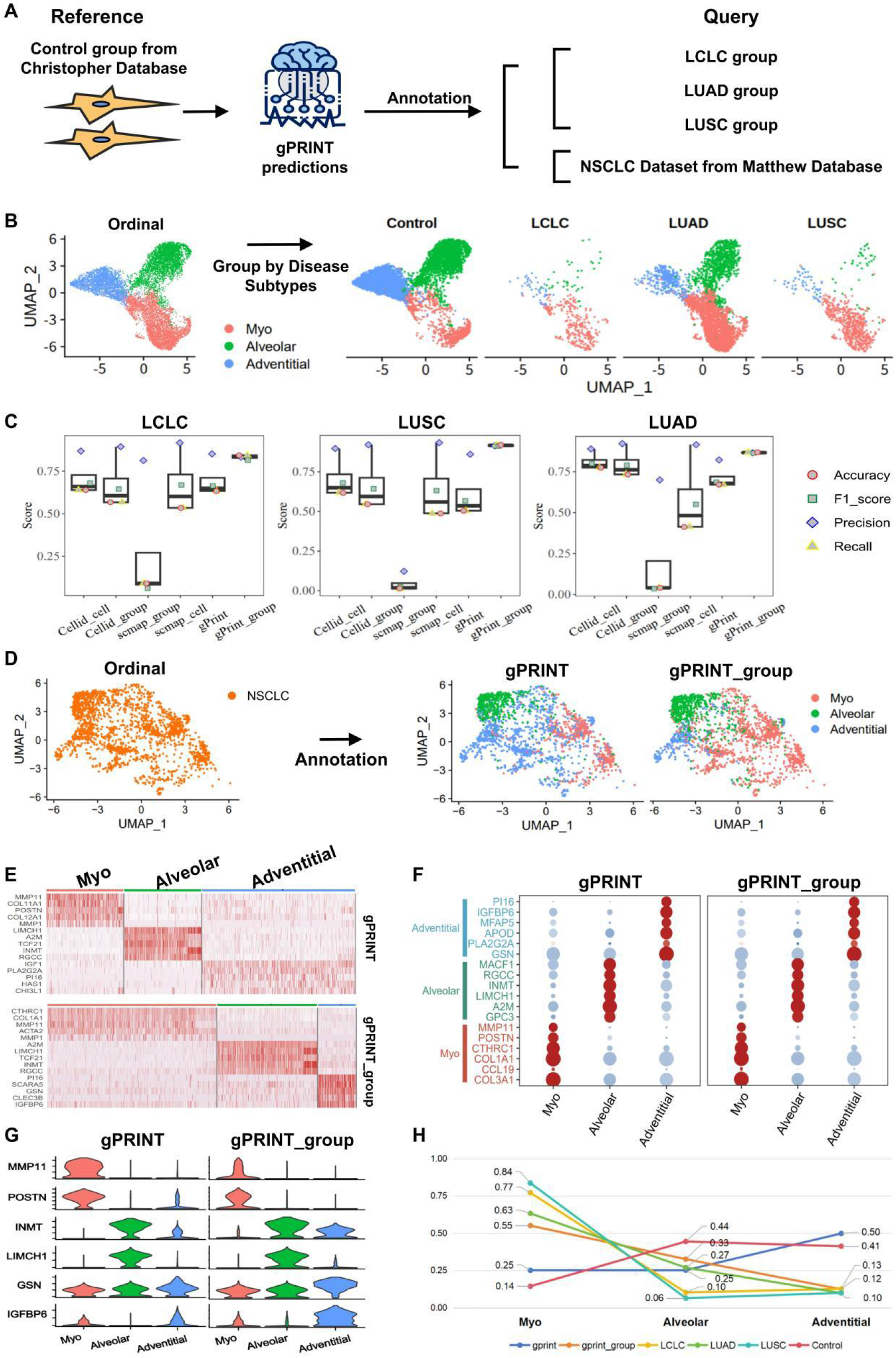
Annotation of refined subtypes on fibroblasts across NSCLC disease subgroups and identification of key disease subpopulations. (A) Demonstration of gPRINT’s cross-NSCLC disease subgroup annotation and cross-dataset annotation scheme on the human fibroblast data set. (B) UMAP diagram of cell subtype distribution of NSCLC disease-related fibroblasts. The color of the cells represents the cell subtype. Type, the left represents the NSCLC fibroblast single cell data set in the original article, and the four on the right represent different samples separated by disease subgroup. (C) Boxplots summarize method’s F1 scores, and show the median (center line), interquartile range (hinges) and 1.5 times the interquartile range (whiskers). (D) UMAP diagram shows the annotation results of fibroblasts from Matthew Database using gPRINT and gPRINT_group annotation methods, where the color of cells represents cell subtypes. The expression of the top 5 differentially expressed markers in different cell subtypes was annotated and presented as a heat map. (F) Dot plot showing the expression of marker genes of different cell subtypes on cell subtypes annotated by different algorithms. (G) Violin plot showing the expression levels of marker genes of all different cell types on cell subtypes obtained by different annotation methods. (H) Line chart of the proportion of cell subtypes mapped under different disease subgroups and annotation methods. Different colors represent different data sets.

Comparing the annotation results of different algorithms on fibroblast data sets in various disease states, we found that the gPRINT_group algorithm performed best.gPRINT_group consistently achieved high accuracy (>84%), recall (>84%), and F1 values (>81%) for all evaluated reference-to-query cell-type assignments. The Cellid_cell algorithm ranked second (accuracy (>61%), recall (>61%), and F1 values (>68%)) (Figure 5C,S7). Furthermore, by observing the percentage line graph of different cell subtypes in NSCLC disease subgroups (Figure 5H), we discovered that the Myofibroblast subtype was more predominant in the diseased group compared to the healthy control group, while the Alveolar cell subtype was more prevalent in the LUAD disease subgroup compared to the LCLC and LUSC subtypes. This indicates that the key cell subtype in NSCLC is Myofibroblasts, while the specific key cell subtype in LUAD is the Alveolar cell subtype. This finding suggests that targeted drug exploration can be conducted based on different cell subtypes for different diseases and subtypes. In summary, the gPRINT algorithm effectively addresses the precise mapping of cell subtypes and enables cell type mapping across different disease states. This contributes to understanding the differentiation and transformation mechanisms of cells under different disease subgroups and their roles in disease occurrence and development, providing guidance for targeted drug development or exploration.

Furthermore, we applied the aforementioned model to annotate a non-small cell lung cancer (NSCLC) fibroblast dataset from the Matthew dataset^[38]^, mapping the fibroblast subtype labels across different datasets (Figure 5D). By observing the differential gene heat map after annotation (Figure 5E), we found that gPRINT and gPRINT_group algorithms successfully annotated three distinct fibroblast subgroups. Additionally, by examining the expression bubble plots and violin plots of marker genes corresponding to the three cell subtypes (Figure 5F,G), we determined that our annotation was generally accurate, although some differences were observed. We then observed the cell proportion line graph of different cell subtypes in the Matthew dataset^[38]^ (Figure 5H) and found that the annotated dataset’s cell subtype proportions closely matched those of the LUAD disease subgroup. Based on this observation, we inferred that the Matthew dataset^[38]^ represented the LUAD disease subgroup, suggesting the need for more precise and tailored treatment strategies for this subtype. Overall, we believe that the gPRINT algorithm can effectively solve the problem of accurate mapping of cell subtype labels across data sets and learn from existing fibroblast atlases to provide a cell type reference for future fibroblast research.

## Discussion

In this study, we introduced the concept of “gene print” expression pattern for individual cells. Utilizing this pattern along with neural network algorithms, we developed the gPRINT tool for uniform annotation of disease-specific cell subtypes. Since genes often exhibit “co-expression” patterns (Figure S2), with possible correlations between adjacent genes, we sorted the gene expression matrix according to HG38. This sorting allowed us to obtain unique one-dimensional “gene print” expression patterns for each cell. Unlike the matrix format, this pattern incorporates positional information among genes, making it more closely resemble the actual gene expression. Within this pattern, by applying principles akin to voice recognition filtering, such as adding convolutional layers, we were able to address batch effects more effectively, similar to noise reduction in voice recognition, ultimately improving the accuracy and reliability of cell type and subtype identification. Therefore, the gPRNIT algorithm, based on neural network models and the data representation of “gene print,” effectively tackles issues of cellular heterogeneity and noise in single-cell RNA sequencing data, enhancing accuracy and reliability in identifying cell types and subtypes.

Specifically, in this study, we first validated the feasibility of our algorithm on internal test sets from pancreatic tissues and the HCL database across various tissue types. We successfully achieved cross-dataset cell type mapping in pancreatic tissues and compared it with other algorithms. The gPRINT algorithm achieved the highest accuracy of 97.39% in annotating high-quality datasets in pancreatic tissues (Figure 2). When annotating low-quality pancreatic datasets, gPRINT achieved an average accuracy of 98.37%, followed by SingleR with an average accuracy of 95.87% (Figure 3). This performance was significantly higher than other cell annotation algorithms. These results indicate that the gPRINT algorithm can achieve the level of existing algorithms, or even surpass them, in cell type annotation at the level of large subpopulations. Overall, the gPRINT algorithm demonstrates its effectiveness and superiority in addressing the challenges of cell heterogeneity and noise in single-cell RNA sequencing data, thereby improving the accuracy and reliability of cell type and subtype identification, as evidenced by its performance in pancreatic tissue annotation across datasets (Figure 3).

Next, we further conducted cross-time cell type mapping on the human skeletal muscle atlas. In the annotation of the human fetal skeletal muscle atlas, the gPRINT and gPRINT_group algorithms successfully mapped cell types and subtypes from the Fetal Wk 12-14 data to the Fetal Wk 17-18 data with high accuracy (>90% and >92%). Following closely behind was the Cellid_cell algorithm, with an accuracy of (>82%). In the adult limb skeletal muscle atlas, the gPRINT and gPRINT_group algorithms also achieved high accuracy (>92% and >93%) in mapping cell types and subtypes from the Postnatal Juvenile Yr7-11 data to the Adult Yr34-42 data. Again, Cellid_cell had an accuracy of (>64%) (Figure 4). These results demonstrate that the gPRINT algorithm can accurately perform cell type and subtype label mapping across different time points and datasets.

Additionally, we conducted precise fibroblast subtype mapping across subtypes of non-small cell lung cancer (NSCLC). The gPRINT algorithm exhibited an average accuracy of 87.67% (Group), significantly outperforming the Cellid (Cell) algorithm with an accuracy of 67.72% (Figure 5). This indicates that the gPRINT algorithm is more precise than other methods in mapping cell subtypes under different disease subgroups. This capability will aid in understanding the composition of cells under different disease subgroups and their roles in disease initiation and progression, providing valuable insights for targeted drug development and exploration.

These research findings demonstrate that gPRINT not only provides a new method and approach for cell type annotation but also exhibits strong reliability and accuracy in cross-dataset and cross-time cell type mapping. Furthermore, the validation of gPRINT algorithm for cell type annotation under different disease states has been achieved. These discoveries hold significant implications for understanding cell type differences in various health and disease states and shedding light on the mechanisms of disease development. Additionally, the methods used in this study can serve as a reference and inspiration for cell type annotation and cross-dataset cell type mapping in other research contexts. In summary, the results of this study offer fresh insights and methods for the analysis and interpretation of single-cell transcriptome data. They are of critical importance for understanding cell type diversity and the mechanisms underlying disease, paving the way for valuable extensions and applications in future cellular and biological research.

However, this study also has certain limitations. Firstly, the choice of reference datasets may impact the accuracy and generalizability of the algorithm: larger sample sizes and more extensive coverage can yield more accurate annotation results. Secondly, the selection of algorithms and parameter settings may affect the stability and reliability of the results. Therefore, in future research, further improvements and refinements of the algorithm can be pursued to enhance the accuracy and efficiency of cell type annotation and cross-dataset cell type mapping. In terms of cell type annotation, further exploration of cell type differences in different tissues and organs, as well as cell type variations associated with disease states, can be conducted. These studies can provide deeper insights into cell type diversity and function, aiding in the diagnosis and treatment of diseases. Moreover, in the realm of cross-dataset cell type mapping, additional datasets can be used for validation to improve the algorithm’s reliability and generalizability. Application of this algorithm to single-cell data from other biological species can increase our understanding of biodiversity. Finally, on the methodological front, further exploration and enhancement of techniques such as data preprocessing, feature selection, and model construction can improve the accuracy and efficiency of cell type annotation and cross-dataset cell type mapping. In conclusion, this study offers a new method and approach for cell type annotation and cross-dataset cell type mapping, while also suggesting valuable directions for future research. Additionally, addressing and resolving technical variations and differences between different datasets can enhance the reliability and generalizability of cross-dataset cell type mapping.

## Methods

### 1. Data Preprocessing

All scRNA-seq data were preprocessed using R (version 3.6.1). For single-cell data, genes with a total expression less than 2 were removed. When a dataset served as a reference set, cell types with fewer than ten cells were excluded because models trained on datasets with too few cells may overfit. This requirement was not applied when the dataset served as a test set. Genes in both the reference and test sets were ordered according to the gene order in HG38. The test set required genes consistent with those in the reference set, and missing genes in the test set were filled with zeros. The Seurat package’s NormalizeData and ScaleData functions were used to standardize and normalize the data in preparation for running the gPRINT method.

If the gPRINT_group algorithm was used, marker genes for each cell type from a marker gene database (e.g., PanglaoDB database^[40]^) or calculated using the Seurat package’s FindAllMarkers function were used. These genes were then converted into one-dimensional “feature pattern” based on the order of the HG38 reference genome and input into the gPRINT_group model.

### 2. Obtaining “gene print”

Just as each person has their unique fingerprint and voiceprint, each cell has its “gene print”. The gPRINT algorithm first rearranges genes according to the end value of the transcript of each gene in the human reference genome sequence HG38, and then draws the waveform of the expression level of each gene in each cell to obtain the “gene print” to which the cell belongs. In the HG38 reference genome, the termination site (end) of the transcript of each gene is usually unique. So we can fix the position information of the “gene print” pattern. This pattern is akin to the amplitude envelopes of soundwaves. Each gene can be seen as a frame segment, with the gene’s expression value resembling the value on the amplitude envelope of a soundwave. Then, this “gene print” is analyzed using deep learning algorithms commonly employed in soundwave recognition. Deep learning has matured significantly in applications such as sound classification, speech recognition, speech-to-text conversion, and interactions with intelligent assistants. Sound and audio are fundamentally time-series data, and when extracting features from soundwaves, amplitude envelope extraction is commonly employed. The purpose of the amplitude envelope is to extract the maximum amplitude for each frame and concatenate them together. Amplitude represents the signal’s volume (or loudness). The signal is first decomposed into its constituent windows, and then the maximum amplitude within each window is identified. These maximum amplitudes are then connected over time to form the amplitude envelope.

In our proposed algorithm, we leverage the principles of deep learning applied in soundwave recognition to treat the positions of open gene expressions as temporal information in soundwaves. Each gene interval is viewed as a frame segment in the soundwave, and scRNA-seq data is transformed into waveform data for learning and annotation. This approach introduces a new feature: gene position information.

As is well-known, DNA is arranged in an ordered sequence on each chromosome (Figure S1). Histones within nucleosomes, as the axis around which DNA is wound, can influence gene transcription through specific translation modifications at certain sites. Additionally, genes that are physically clustered on the same nucleosome may be co-regulated and exhibit similar expression patterns, resulting in “co-expression”. Therefore, adding positional information is meaningful for each cell. We incorporate this feature into the algorithm, reshaping the gene expression patterns into one-dimensional “gene print”. This addition of a new feature, which closely resembles the form of gene expression, theoretically contributes to the improvement of annotation precision in cell annotation.

### 3. Construction of Neural Network Methods for Single-Cell Annotation

Based on the “gene prints”, we use a one-dimensional convolutional neural network (1D CNN), which comprises an input layer, five hidden layers, and an output layer. The input layer has the same number of nodes as the reference genes. The hidden layers consist of 2 convolutional layers, 1 pooling layer, 1 Flatten layer, and 1 fully connected layer. The pooling layer is used to retain essential features while reducing computation load, the Flatten layer connects the convolutional layers with the Dense fully connected layer. The primary role of the fully connected layer is to apply nonlinear transformations to the features extracted by the preceding convolutional layers, extracting correlations between these features, and mapping them to the output space. The input dimension of the model is based on the number of genes in the reference dataset, the convolutional layers have 16 kernels with a kernel size of 3, and the Dense layer uses the softmax function.

Based on the “feature patterns”, we use a backpropagation (BP) neural network. The BP neural network is a multi-layer feedforward neural network that consists of two main stages in its training process: forward propagation of signals and backward propagation of errors. It employs the steepest descent method as the learning rule, adjusting the network’s weights and thresholds continuously through backpropagation to minimize the sum of squared errors.

### 4. Evaluation Metrics

When performing classification tasks in machine learning, common evaluation metrics for model performance include precision, recall, accuracy, F1-score, and receiver operating characteristic (ROC) curve. Precision refers to the proportion of true positive samples among all samples predicted as positive. Recall, also known as sensitivity, represents the proportion of true positive samples among all actual positive samples. Accuracy is a performance metric for classification tasks, indicating the proportion of correctly classified samples out of all samples. F1-score, also called the balanced F-score, is the harmonic mean of precision and recall, ranging from 0 to 1, where a higher value indicates better classification results. The receiver operating characteristic (ROC) curve is plotted with the false positive rate (FPR) on the x-axis and the true positive rate (TPR) on the y-axis, and the area under the ROC curve (AUC) provides a comprehensive evaluation of the model’s classification performance. AUC ranges from 0 to 1, with higher values indicating better classification performance.

For each cell type in each test dataset, the following formulas are used to calculate precision, recall, accuracy, F1-score, and ROC curve to evaluate the model:

Precision = TP / (TP + FP)

Recall = TP / (TP + FN)

Accuracy = (TP + TN) / (TP + TN + FP + FN)

F1-score = 2TP / (2TP + FP + FN)

TPR = TP / (TP + FN)

FPR = FP / (FP + TN)

Where:

TP: True Positive, correctly predicted positive samples.

FP: False Positive, incorrectly predicted positive samples.

TN: True Negative, correctly predicted negative samples.

FN: False Negative, incorrectly predicted negative samples.

## Data Availability

Our laboratory has previously accumulated multiple validated single-cell transcriptome datasets, which are available for model validation and application testing. Additionally, we can access single-cell transcriptome and epigenome data from various tissues and organs through the Gene Expression Omnibus (GEO) database. Specifically, datasets from Wang^[20]^, Xin^[21]^, and Muraro^[22]^ for pancreatic scRNA-seq, Xi^[31]^ for muscle and tendon-related data, Li^[25]^ for human colorectal tumor data, Camp^[23]^ for human brain tissue-related single-cell data, human non-small cell lung cancer (NSCLC) disease subgroup maps^[32]^, and their associated cell type annotations were downloaded from GEO (GSE83139, GSE81608, GSE85241, GSE147457, GSE81861, GSE75140, and GSE153935). The dataset from Segerstolpe^[24]^ was obtained from ArrayExpress (EBI) (E-MTAB-5061). Notably, the Wang and Xin datasets used the SMARTer method for single-cell RNA sequencing, the Muraro dataset utilized CEL-Seq2, and the Segerstolpe dataset was generated using Smart-Seq2 sequencing. For human perturbed state fibroblast atlas data^[38]^, it can be accessed through https://fibroXplorer.com. The human pancreatic cancer single-cell data is available in the EGA database^[6]^ (EGAD00001005365). Many single-cell datasets from various tissues are sourced from the Human Cell Landscape (HCL) database^[7]^: http://bis.zju.edu.cn/HCL/.

## Acknowledgments

This work was supported by the NSFC grants (T2121004, 82072463, 32271406, 82222044, 82202045),the National key research and development program of China (2021YFA1100500), Zhejiang Provincial Natural Science Foundation of China (LR20H060001, LZ22H060002), Fundamental Research Funds for the Central Universities, the General Research Fund of the Research Grants Council of Hong Kong (24101921).

