## Supplementary material for "Cross-Dataset Identification of Human Disease-Specific Cell Subtypes Enabled by the Gene Print-based Algorithm--gPRINT": Supplementary material2.pdf

Figure S1. Algorithm principle diagram and schematic diagram.

Figure S2. Confusion matrix for internal test set.

Figure S3. nCount_RNA values for each data set of pancreatic tissue.

Figure S4. Annotations of different algorithms for Wk12_14 and Wk17_18 data sets.

Figure S5. Confusion matrix of human fetal skeletal muscle across data sets.

Figure S6. Annotation application across time on adult skeletal muscle atlas.

Figure S7. Beeswarm plot display of small subpopulation annotation evaluation indicators under fibroblasts across NSCLC disease subtypes.

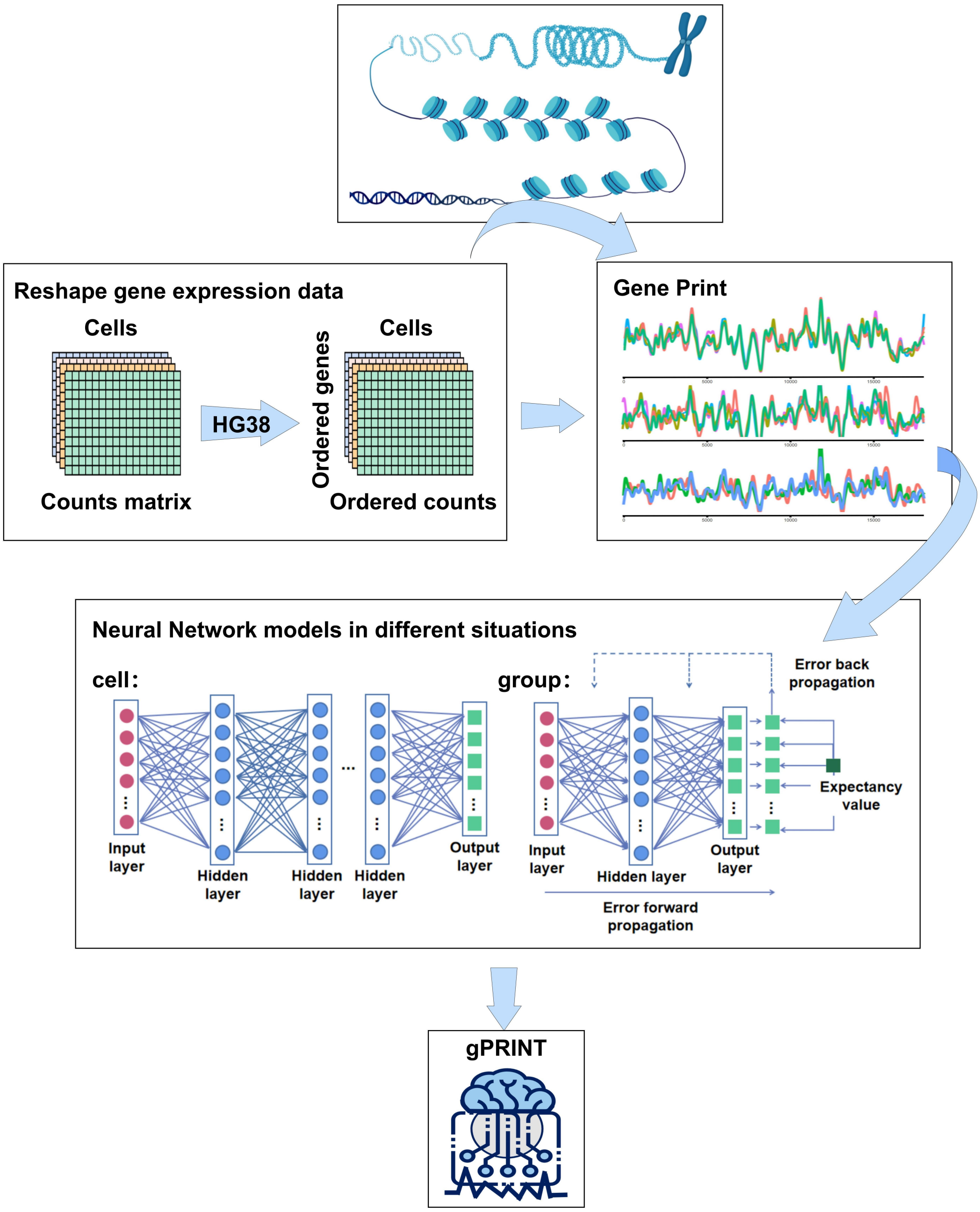


Figure S1. Algorithm principle diagram and schematic diagram.

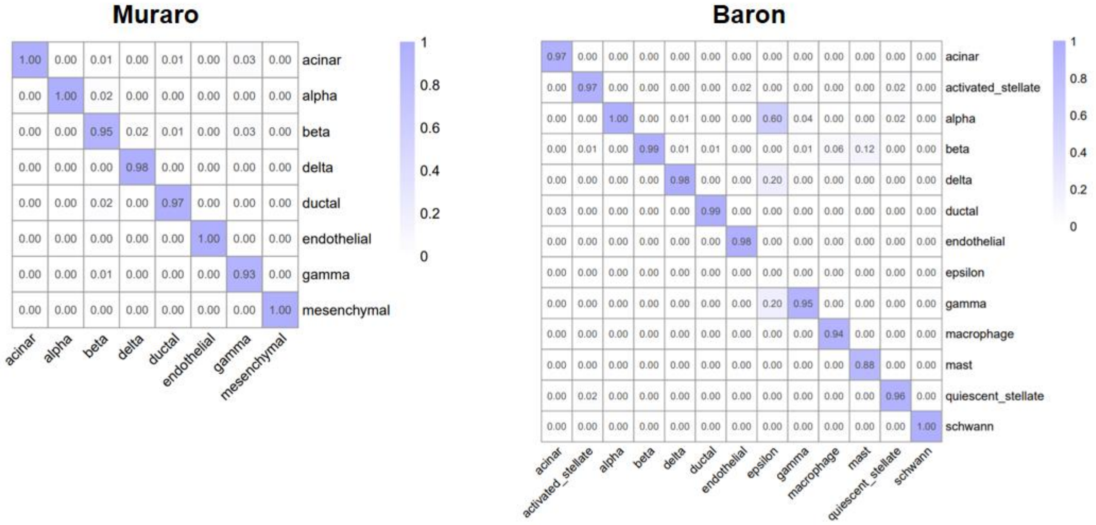


Figure S2. Confusion matrix for internal test set.

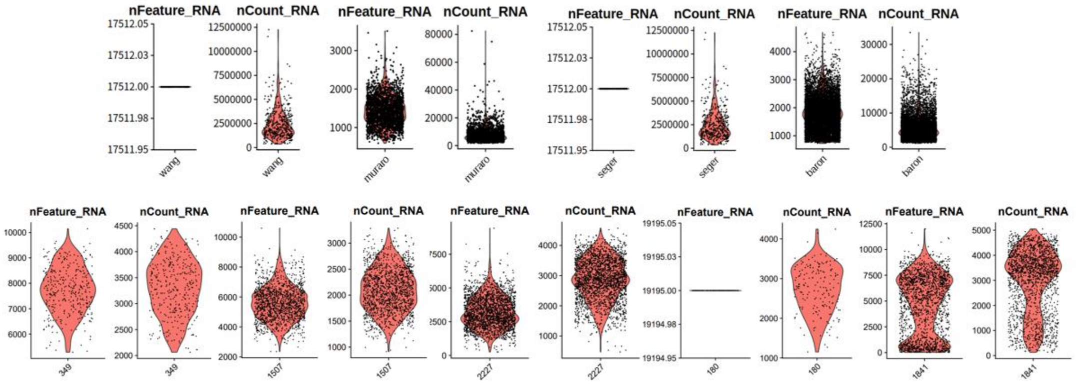


Figure S3. nCount_RNA values for each data set of pancreatic tissue.

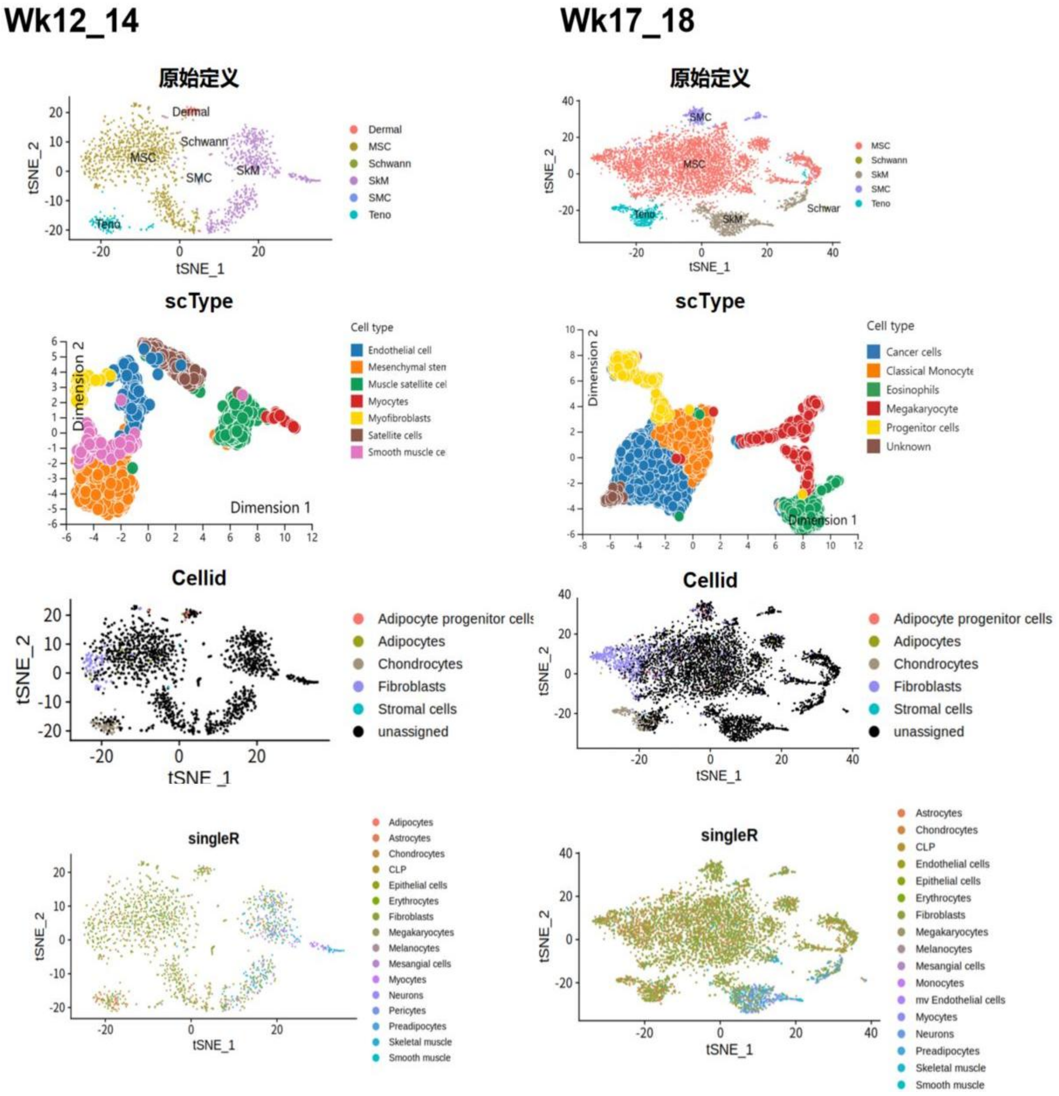


Figure S4. Annotations of different algorithms for Wk12_14 and Wk17_18 data sets.

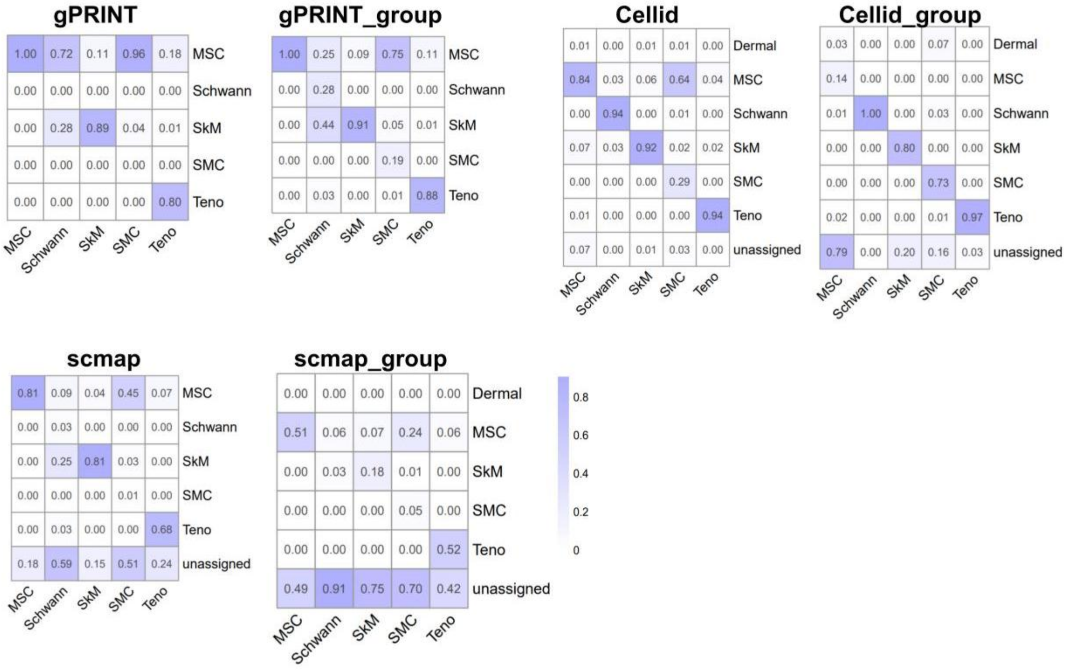


Figure S5. Confusion matrix of human fetal skeletal muscle across data sets.

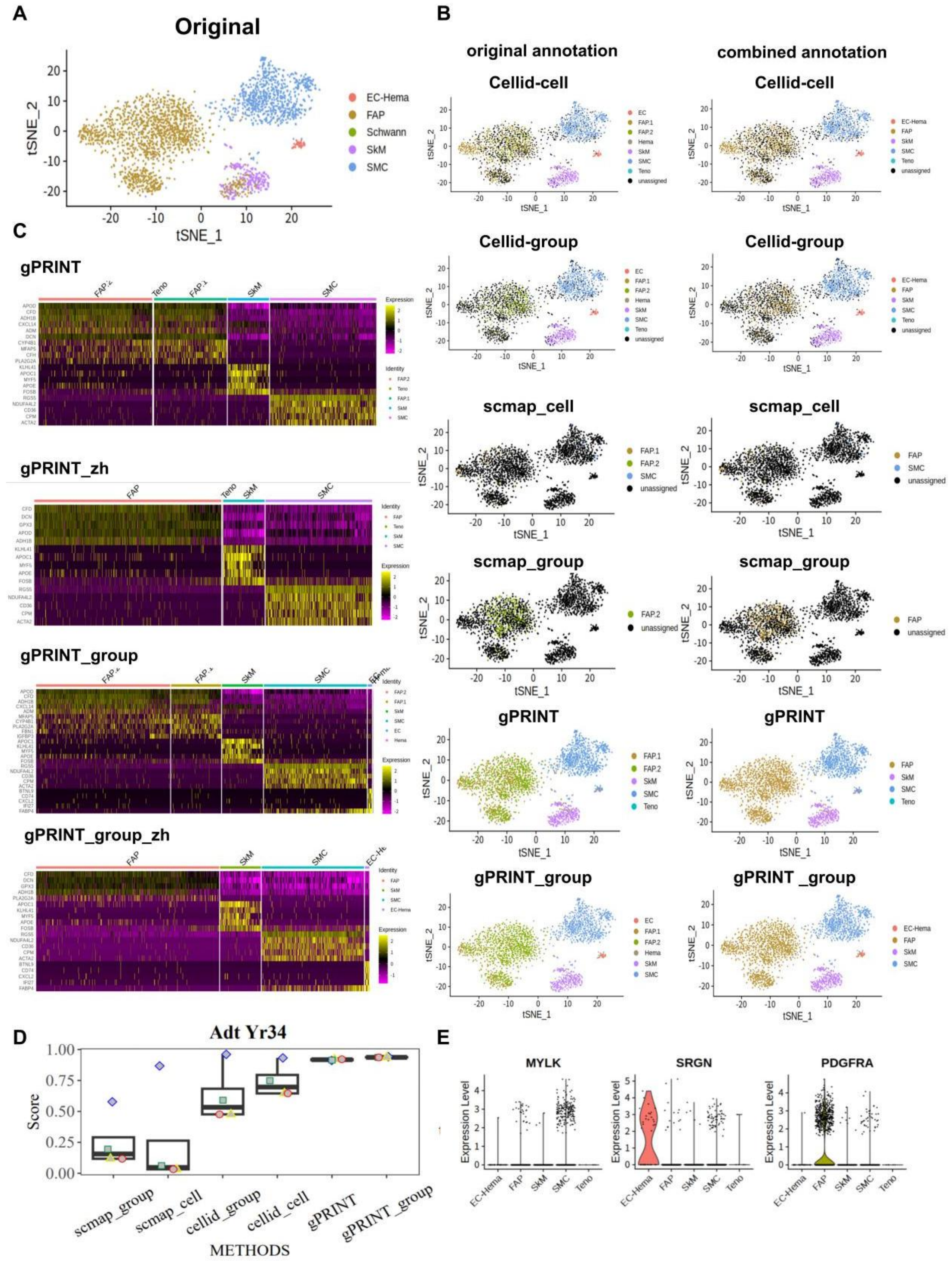


Figure S6. Annotation application across time on adult skeletal muscle atlas. (A) Cell subtype mapping across tendinopathy and healthy groups in humans. Using the single cell data of the healthy group as a reference, the identity mapping of cell subtypes on the tendinopathy data set was completed. (B) Confusion matrix representations comparing manual annotations with the classification labels generated by gPRINT, gPRINT_group, Cellid and Cellid_group. Diagonal represents percentage of labels that agree between both classification methods. (C) Percentage of mapped cell subtypes in gPRINT mode Column chart. Yellow is healthy, blue is tendinopathy. (D) The process of predicting key target genes and screening related drugs based on the found disease-specific cell subtypes.

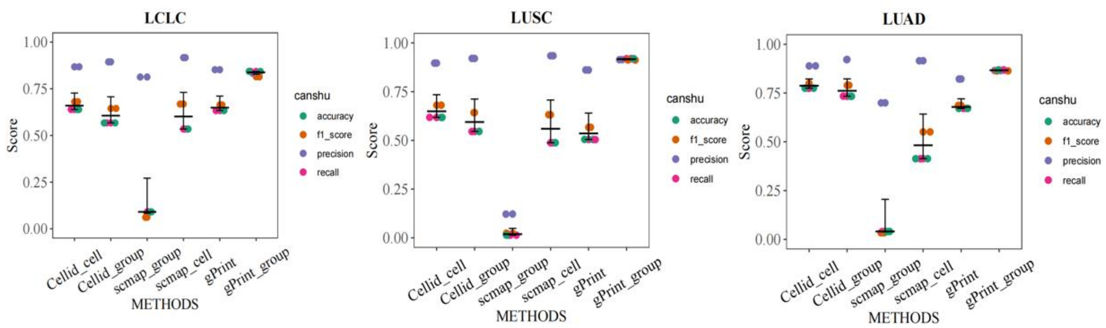


Figure S7. Beeswarm plot display of small subpopulation annotation evaluation indicators under fibroblasts across NSCLC disease subtypes.
